# baerhunter: An *R* package for the discovery and analysis of expressed non-coding regions in bacterial RNA-seq data

**DOI:** 10.1101/612937

**Authors:** A. Ozuna, D. Liberto, R. M. Joyce, K.B. Arnvig, I. Nobeli

## Abstract

**Summary:** Standard bioinformatics pipelines for the analysis of bacterial transcriptomic data commonly ignore non-coding but functional elements e.g. small RNAs, long antisense RNAs or untranslated regions (UTRs) of mRNA transcripts. The root of this problem is the use of incomplete genome annotation files. Here, we present *baerhunter*, a method implemented in *R*, that automates the discovery of expressed non-coding RNAs and UTRs from RNA-seq reads mapped to a reference genome. The core algorithm is part of a pipeline that facilitates downstream analysis of both coding and non-coding features. The method is simple, easy to extend and customize and, in limited tests with simulated and real data, compares favourably against the currently most popular alternative.

**Availability:** The *baerhunter R* package is available from: https://github.com/irilenia/baerhunter

## Introduction

Next-generation sequencing has facilitated global surveys of the transcriptome, largely focused on studying differential expression of genes across different conditions. Studies of eukaryotic transcriptomes are increasingly embracing the analysis of non-coding transcript expression but in bacteria, where intergenic regions tend to be a lot shorter (Thorpe *et al.*, 2017) and less well annotated, automated differential gene expression is still largely synonymous with differential expression of the coding regions (CDS). As the functional importance of bacterial non-coding RNAs (ncRNAs - the term used here to cover long antisense RNA, small regulatory RNA (sRNA), and untranslated parts of mRNAs) is becoming evident (Michaux *et al.*, 2014), so is the need for including them in differential expression studies.

A major obstacle in studying non-coding RNA expression in bacteria is that relatively few ncRNAs are reliably annotated and, with the exception of well-known cases (such as tRNAs, ribosomal RNAs and, more recently, some members of the RFAM (Kalvari *et al.*, 2018) families), the majority are not included in the standard annotation files required by computational pipelines. Requiring the non-coding RNAs to be included in the annotation is prohibiting their analysis by methods such as TrBorderEx (Wang *et al.*, 2015), which identifies the transcript boundaries but does not find new non-coding RNAs. An alternative to waiting for annotations to improve is to identify ncRNAs using the expression data signal. Early efforts in this direction relied on a combination of manual inspection and in-house written scripts to identify clusters of reads falling outside known CDS regions (Arnvig *et al.*, 2011)(Wilms *et al.*, 2012)(Pfeifer-Sancar *et al.*, 2013). These studies offered great insights into the non-coding transcriptome but applying their approach in a different context is time-consuming and prone to errors due to the need for recreating the pipelines from scratch. Selected studies have led to publicly available software for the study of ncRNAs in bacteria. However, some methods are limited to specific species (Pellin *et al.*, 2012) or rely on specialized sequencing protocols (Peña-Castillo *et al.*, 2015; Amman *et al.*, 2014). Two notable exceptions have appeared in recent years. DETR’PROK (Toffano-Nioche *et al.*, 2013) employs the Galaxy platform (Afgan *et al.*, 2018) to classify clusters of RNA-seq reads not overlapping with annotated genes as sRNAs, antisense RNAs, and UTRs. The updated annotation file can be used to carry out differential gene expression. However, the DETR’PROK workflow is composed of a large number of steps, requires an active user input at several stages and depends on access to a Galaxy instance. Rockhopper (McClure *et al.*, 2013), a standalone, Java-based program that allows both identification of features in a bacterial transcriptome and differential expression between conditions, is primarily aimed at non-bioinformaticians. A user-friendly graphical interface masks a fairly sophisticated set of algorithms that are presented as a black box with only a handful of parameters accessible to the user. Although straightforward to use, the set up is inflexible with little scope for extending or altering the pipeline without expert interfering with the code.

Here, we present *baerhunter* (“baer” stands for **ba**cterial **e**xpressed **r**egions), a new method implemented in *R* (R Core Team, 2018), for automating the detection and quantification of expressed putative non-coding RNAs (including UTRs) in bacterial strand-specific RNA-seq data. At the core of the *baerhunter* pipeline is a simple but effective method of capturing expressed intergenic regions across sets of RNA-seq data samples. The method is designed to provide predictions of approximate locations of non-coding elements, reflecting our belief that accurate definitions of transcript ends are best achieved by targeted experimental methods rather than computational predictions from noisy data. The pipeline built around this method facilitates the analysis of differential expression of these regions in parallel with the more traditional protein-coding-focused analysis. Below, we describe our method and present the results of testing its performance both on simulated and real data from *Mycobacterium tuberculosis* (*Mtb*). In addition, we compare *baerhunter* to Rockhopper, chosen as the most widely used alternative method.

## Methods

The core algorithm of *baerhunter* carries out a search for intergenic features on each strand, displaying a minimum length and coverage depth in the RNA-seq signal (see Supplementary Methods for details). The algorithm is wrapped within a “driver” *R* script that can be easily edited to include, exclude or modify steps, depending on the user’s requirements. Individual functions of *baerhunter* can also be used in isolation or incorporated within different pipelines. In the default mode, *baerhunter* reads in a set of Binary Alignment Map (BAM) files with RNA-seq reads mapped to a reference genome and an annotation file in the Generic Feature Format (GFF3) for the same genome. It will then identify expressed intergenic regions on each strand (“features”) and combine overlapping features across multiple BAM files to create a full set of non-overlapping genomic features. Features are classified as either “UTR”, if they are thought to be the untranslated part of a coding mRNA or “sRNA” (used in this context to encompass all other types of non-coding RNA in bacteria, including long antisense RNAs). In addition, *baerhunter* allows for new transcripts to be filtered by their expression level (normalised to transcripts per million (TPM) values), as many very low-expression features are likely to be the result of transcriptional noise or ambiguous read mapping. Finally, differential expression analysis, including all newly annotated putative features, is facilitated by a wrapper script that utilizes the DESeq2 method (Love *et al.*, 2014).

To test *baerhunter*, a simulated RNA-seq dataset was created using the package *polyester* (Frazee *et al.*, 2015). In addition, RNA-seq data from the study of (Cortes *et al.*, 2013), six samples from exponentially growing and starved cultures of *Mtb*, were downloaded from Array Express (E-MTAB-1616) and processed as detailed in Supplementary Methods. Following analysis with *baerhunter*, the sRNA/UTR predictions were compared to a set of experimentally confirmed and predicted mycobacterial sRNAs from the comprehensive review of (Haning *et al.*, 2014). Transcription start sites reported by (Cortes *et al.*, 2013) for the same samples were also used to assess the accuracy of our predictions. The genome browser Artemis (version 17.0.1) (Carver *et al.*, 2012) was used for visualization.

Rockhopper was used with default parameters, except for the minimum transcript length that was set to 40 nucleotides to match the *baerhunter* settings. Two minimum expression thresholds were tested (0.5, the default, and 0.2, to increase sensitivity).

All code and data required to reproduce this analysis are available from the github repository: https://github.com/irilenia/baerhunter_paper

The latest version of baerhunter is available from: https://github.com/irilenia/baerhunter

## Results

### Simulated dataset

We tested the ability of *baerhunter* to recover expressed intergenic regions and UTRs using simulated data. 1000 genomic features were randomly selected from the *Mtb* genome, including twenty-four short RNAs included in the original annotation (see Supplemental Methods). As the genome annotation file does not include UTR information for *Mtb*, artificial UTRs were added to a random subset of 200 genes. These 1000 features were simulated in 10 samples belonging to two groups (with fold changes between 1 and 5 applied to 20% of the features). Our pipeline applied to paired-end read simulations recovered all short RNAs and all UTRs, with exact predictions for the start and end coordinates of all 24 sRNAs and over half of the UTRs (the remaining being in their vast majority within 5 nucleotides of the true range). Results were relatively insensitive to small changes in the program parameters (Supp Table 1). Rockhopper performed similarly on sRNAs, recovering 23 of 24 when run at default sensitivity (22 of the 23 sRNAs had their coordinates exactly predicted) but was less successful in the prediction of UTRs, missing 4 of the 200 and estimating the lengths of approximately a quarter of the ones it predicted to be at least 20 nucleotides shorter than expected (Supp Figure 4).

### Real dataset

In addition to using simulated data, we applied *baerhunter* to the RNA-seq data from starved and exponentially grown cultures of *Mtb* (Cortes *et al.*, 2013). In this case, the true number of non-coding RNAs is unknown, so *baerhunter* was benchmarked against transcription start-site data from the same samples, as well as lists of known and predicted non-coding RNAs (Haning *et al.*, 2014) in order to assess the likely accuracy of our predictions.

At the more stringent parameter values (5-20), 74-83% of the predicted sRNA features in samples from either condition are supported by the presence of a TSS, even at one-nucleotide resolution (Supp Figure 5A & Supp Table 2). Relaxation of the cut-off (to 5-10) increases false positives but, importantly it also increases true predictions, thus allowing more transcripts to be discovered at the cost of a more noisy output (Supp Fig. 5B & Supp Table 2). Although more than half of the *baerhunter*-predicted sRNAs do not correspond to sRNAs in the published list of (Haning *et al.*, 2014), visual examination of the RNA-seq signal confirms expression at these loci (Supp Figure 6), usually from a very weak TSS that has not passed the inclusion cut-off in the original study by (Cortes *et al.*, 2013). These transcripts are often expressed at very low levels and can be easily filtered out using expression strength. Rockhopper, run at default expression cut-offs, not only predicts fewer sRNAs but also a smaller percentage of these predictions (∼50-75%) are supported by TSS evidence (Supp Figure 5C&D). The prediction of UTRs is harder to assess. In the absence of ground truth for 3’ UTRs, we compared the start of the 5’ UTR predictions to TSS data. Rockhopper retrieved more of the 5’ UTRs using default sensitivity but differences were generally small (Supp Figure 7). Importantly, the two programs differ in their treatment of 5’ UTRs that are followed by a significant drop near the start of the CDS. Rockhopper annotates these as independent sRNAs, a design that occasionally leads to the unintended consequence of predicting a much shorter 5’ UTR with no experimental support (Figure 1A). In *baerhunter*, we prefer to treat these as 5’ UTRs, acknowledging that they originate from mRNAs containing transcription-terminating regulatory elements ahead of the coding region.

**Figure 1.**
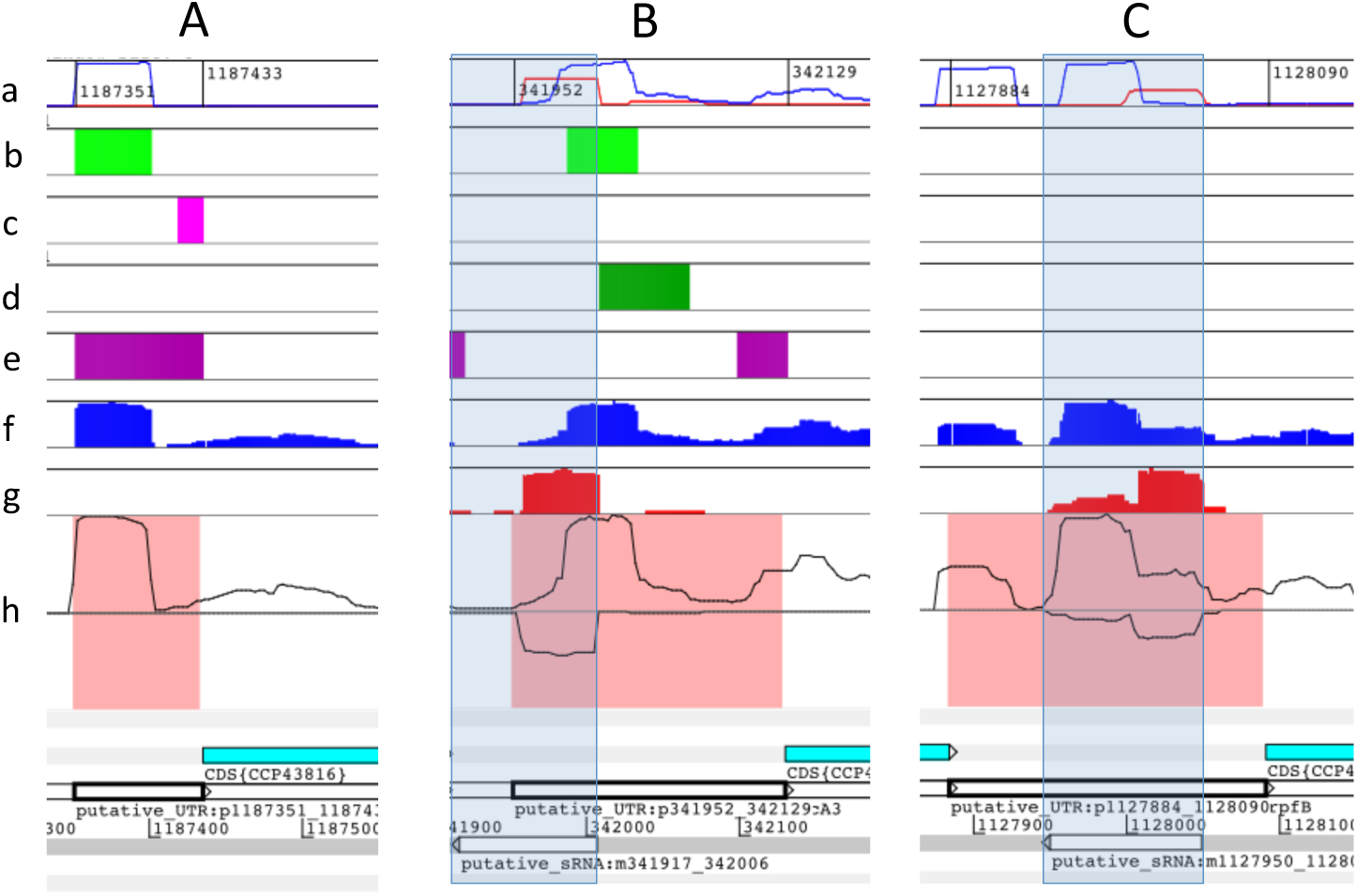
Baerhunter’s predictions of UTRs and small RNAs are supported by experimentally detected transcription start sites (TSS) In all three figures (A, B and C), the panels arranged in rows are: (a) experimentally detected TSS (normalised counts per base from the *Cortes et al.* (Cortes *et al.*, 2013)dataset); (b) and (c), sRNA and UTR predictions from Rockhopper run with default sensitivity, 0.5; (d) and (e), sRNA and UTR predictions from Rockhopper run wih increased sensitivity, 0.2; (f) and (g) RNA-seq trace from Rockhopper mapping of reads for sample ERR262980 (blue: positive strand, red: negative strand); and (h) the RNA-seq trace corresponding to read coverage in sample ERR262980 mapped by our own pipeline (see supplemental Methods). Filled pink rectangles in row (h) highlight the putative UTR region as predicted by *baerhunter*. Transparent blue rectangles (panels B and C) highlight the two short RNAs predicted by *baerhunter* on the negative strand. **A.** The experimentally detected TSS (blue line in row a) supports a 90 nt 5’ UTR for the uncharacterized protein Rv1065. The *baerhunter*’s prediction is 84 nt long (region highlighted with pink in row h), closely following the RNA-seq trace. Rockhopper’s prediction is similar when run with the more sensitive detection threshold of 0.2 (dark pink rectangle, row e) but run at the default 0.5, it splits the prediction to a “non-coding RNA” (green rectangle, row b) and a much shorter 5’ UTR (bright pink, row c) that is not supported by TSS data. **B.** The *baerhunter* program predicts a long (178 nt) 5’ UTR ahead of the Rv0282 gene and an antisense RNA (90nt) partially overlapping this UTR. Both predictions are supported by experimentally detected TSS (blue and red lines on the positive and negative strand respectively; row a). Rockhopper, run with default precision parameters, predicts a non-coding RNA (light green; panel b) and no 5’ UTR (row c), whereas when run at higher sensitivity, it predicts a non-coding RNA further downstream (dark green, row d) as well as a short 5’ UTR (dark pink, row e), corresponding to a weaker TSS just ahead of the coding region of the gene (small blue peak, row a). In both cases, Rockhopper misses the antisense RNA that is clearly seen in the RNA-seq trace of the exponentially growing bacteria (black trace on the negative strand, row h and red-fill trace, row g). **C.** The *baerhunter* program discovers both the 5’UTR ahead of the Rv1009 (*rpfB*) gene and the antisense RNA overlapping it on the negative strand (both of which have experimental TSS support). Rockhopper does not predict any ncRNA features in the same region, although its own mapping of reads results in clear expression on both strands (rows f and g). The similarities between read coverage as reported by Rockhopper (rows f and g) and our own pipeline (row h) indicate that differences between the Rockhopper and *baerhunter* predictions are not due to differences in the way the reads were mapped to the reference genome but instead are due to the different ways the programs identify expressed regions.

## Conclusion

Our new method, *baerhunter*, allows the extraction of bacterial putative non-coding expressed regions directly from RNA-seq data and facilitates the integration of differential expression studies of coding and non-coding elements in bacterial transcriptomes. Importantly, *baerhunter* compares favourably with the most popular alternative method in tests with both simulated and real data.

