## Supplemental Materials for "baerhunter: An *R* package for the discovery and analysis of expressed non-coding regions in bacterial RNA-seq data"

### Supplementary Methods

#### The *baerhunter* algorithm and workflow

The *baerhunter* package detects expressed intergenic areas, starting from a file of RNA-seq reads mapped to genomic coordinates (BAM file) and following the workflow depicted in Supplementary Figure 1. Processing starts with *baerhunter* loading a BAM file using the *GenomicAlignments R* package (Lawrence *et al.*, 2013). The program can distinguish between paired-end and single-end reads, stranded and reversely stranded libraries; unstranded libraries are not supported because they do not allow antisense RNAs to be detected. The aligned reads are used to construct a coverage vector for each strand by counting the number of reads over each position. The coverage vector represents a series of consecutive genomic intervals with constant coverage. *Baerhunter* scans each coverage vector for genomic intervals with a given minimum number of reads aligned to them, defining expression signal peaks. Subsequently, the detected peaks are filtered by a “minimum width” and a “high-coverage threshold” (these are the main user-defined parameters for *baerhunter*). Then, the genomic coordinates of the selected peaks are extracted and combined with the next peak set using the *IRanges R* package (Lawrence *et al.*, 2013). The process is repeated for all BAM files in the directory, resulting in a complete set of expression peak coordinates.

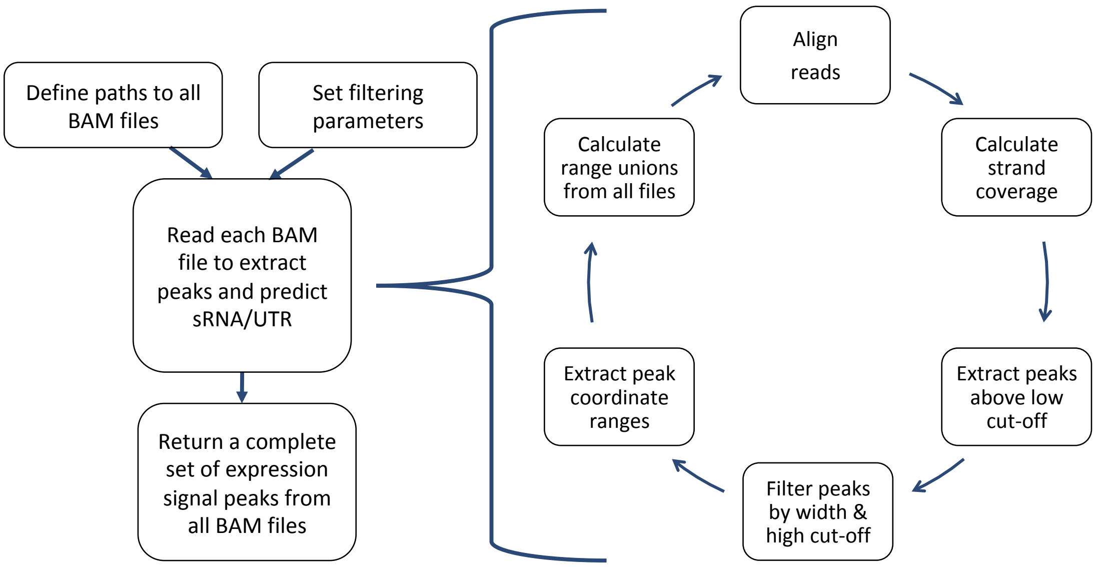


**Supplementary Figure 1. Analysis workflow using *baerhunter* for a single BAM file**
The input file is used to detect expression peaks that satisfy set parameters. The same is repeated for all BAM files in the directory, adjusting the peak coordinate set to include all detected ranges.

Peak detection using two coverage threshold values allows the user to vary the sharpness and borders of the selected expression signal peaks. The low cut-off value is used to separate the peaks from the background signal noise and set the boundaries of each transcript; this is followed by the filtering step that selects only the peaks that contain a consecutive stretch of a given width, which displays coverage values above the high threshold (Supplementary Figure 2A).

Once the peak selection is completed, the genomic coordinates of the detected transcripts are defined. Since the program scans all BAM files in the directory, different transcript sets need to be combined without duplicating transcript information. This is achieved by calculating the transcript boundary union for each strand (Supplementary Figure 2B): the genomic coordinates of the overlapping transcripts are rewritten to include all overlapping features, while all non-overlapping features are recorded as separate transcripts. This allows the capture of transcripts expressed in different conditions without repeating the information obtained from replicate samples. Additionally, the correction of the genomic coordinates achieved by transcript boundary union compensates for potential differences in the transcript lengths caused by technical rather than biological variations, which is beneficial for further processing like differential expression analysis.

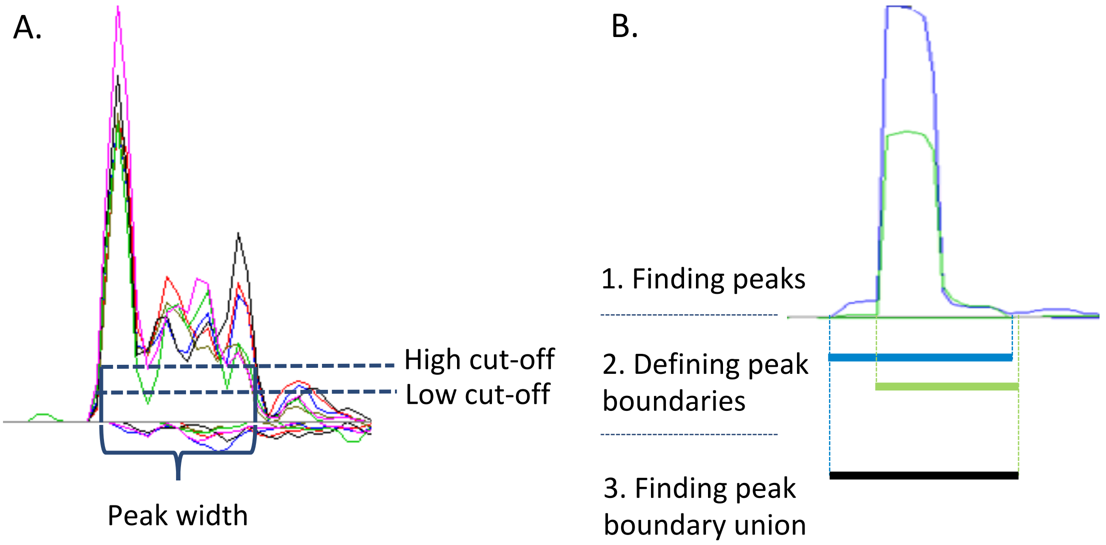


**Supplementary Figure 2.** Peak detection and feature annotation in *baerhunter*.
A) For each RNA-seq trace originating from a sample, three parameters are used to filter the peaks: two are coverage filters and the third is a filter for the peak width. B) Once boundaries are defined for all peaks in each sample, the union of peaks with overlapping coordinates across samples is taken to represent a single feature that is then added to the annotation file.

Once the final transcript set is obtained, the information on the genomic features is extracted from the GFF3 file and used to separate transcripts located in intergenic regions from the ones within annotated genes. It is recommended that the small RNAs present in the original annotation be removed prior to the filtering, allowing the program to redefine such elements according to their expression signal. The transcripts that are located entirely outside the annotated genes are considered as putative sRNAs, while the transcripts that partially span the genes are subjected to further investigation: the parts of the transcripts that do not overlap the genes are isolated and filtered by the given minimum UTR length. The selected features are then annotated as "putative sRNAs" and "putative UTRs". The name assigned to new features comprises information on its genomic coordinates and strand, for instance, sRNA m100_150 is located on the minus strand from position 100 to 150.

The sRNAs and UTRs defined for each strand are then combined with the previously annotated features to produce an ordered strand-specific annotation in accordance with the GFF3 file standards. The attributes column of the predicted elements contains their names and IDs of upstream and downstream features. The surrounding elements are determined assuming that the bacterial chromosomes are circular. The strand-specific annotations are then combined to produce the complete chromosome annotation, which is written as a GFF3 file.

The steps described above can be executed one by one but for ease of use we provide a wrapper function (*feature_file_editor*). This function combines all stages, from BAM file analysis to the annotation of intergenic expressed regions, to produce a complete GFF3 file (Supplementary Figure 3).

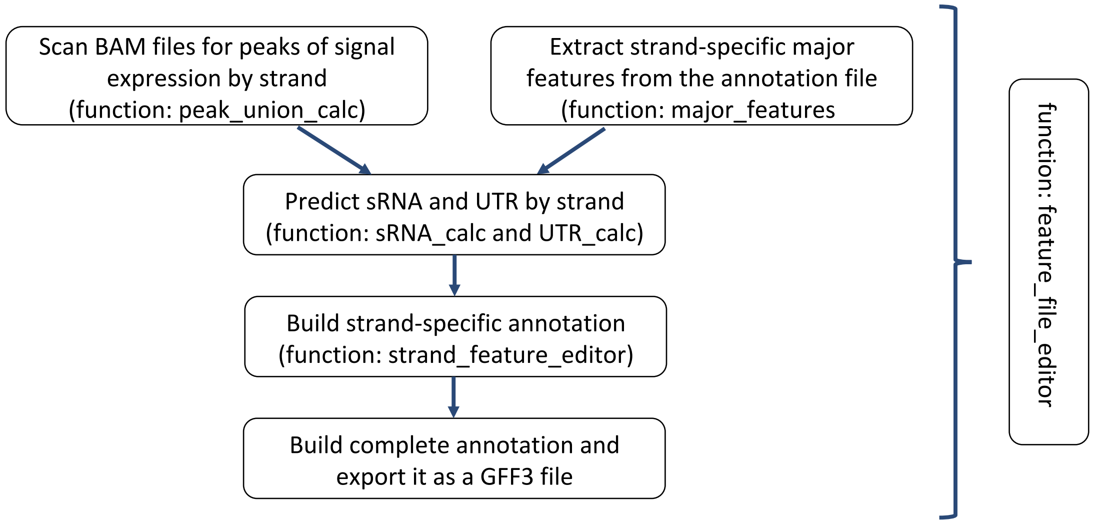


**Supplementary Figure 3.** A workflow of the module of the package responsible for the detection and annotation of intergenic features.

#### Quantification of transcript abundance

The populated GFF3 file can be used for downstream analysis, such as differential gene expression. To allow integration of such analysis, *baerhunter* includes a wrapper function that employs the Rsubread package (Liao *et al.*, 2019) to allocate and quantify the mapped sequencing reads using the newly-produced annotation file. The function parses each major feature type separately, i.e. genes, putative sRNAs and UTRs, and produces distinct count tables.

#### Count normalisation and GFF3 file processing

The detection of expressed regions is done using raw reads, so no count normalisation is applied prior to the prediction of sRNAs and UTRs. The justification for not normalising ahead of the peak detection is that, in our experience, sequencing depth affects the level of "noise" in the data much less than it does the expression of transcripts. Therefore, the initial prediction does not account for the coverage depth differences between replicate samples. However, it is possible to include an estimation of noise based on the actual data for each sample and this is planned for a future version of *baerhunter*.

While over-predicting features might be beneficial in situations where false positives are less detrimental to the analysis than false negatives, *baerhunter* suggests a post-processing step to remove from the new GFF3 file transcripts whose expression is likely to be the result of transcriptional or technical noise or which are expressed at such low levels that are unlikely to have a major biological impact under given conditions. Reads aligned to the features are counted using the Rsubread package, followed by a calculation of Transcripts Per Million (TPM) values for each expressed feature. The TPM values for each feature in a sample should be interpreted as the number of this feature's transcripts that would be observed for every million of RNA molecules sequenced. TPM was selected over the traditional F/RPKM measures (fragments/reads per kilobase per million reads), as it has been argued that TPM values are more consistent and can be, in theory at least, used to compare the proportions of each transcript across different samples, since the sum of the TPMs is equal in each sample

(Wagner *et al.*, 2012). These TPM values can then be used to create flags for the putative sRNA and UTR annotation in the GFF3 file. Flags are added to the attribute column of the file. The flags are set depending on the TPM values across all samples, following the TPM expression level boundaries obtained from the EBI Expression Atlas (https://www.ebi.ac.uk/gxa/home). The rules followed by *baerhunter* are as follows:

- _if TPM values for a feature across all samples are below 0.5, the flag "expression_below_cutoff" is added to the feature;
- if at least one of the TPM values is between 0.5 and 10, the feature is flagged as a "low_expression_hit";
- if any of the TPM values is in the range from 10 to 1000, the feature is flagged as a "medium_expression_hit";
- if any of the values is above 1000, it is considered to be a "high_expression_hit".

Following annotation with expression flags, the user has an option to filter the features by their expression level.

##### Testing for differential gene expression

The detection of differentially expressed genes is performed using the DESeq2 package (Love *et al.*, 2014). The wrapper function provided by *baerhunter*, *differential_expression()*, combines the preparation of the metadata tables with the differential analysis of the count data. The analysis is done in accordance with the standard DESeq2 workflow. The function is able to interpret multiple experimental conditions and incorporate them into the model, although it requires the user to identify the primary condition of interest. The user also needs to provide the cut-off value that is used to filter the count tables: only the features with read counts above the given threshold in at least half the samples will be kept for further analysis. The results of the differential expression analysis and the summary statistics are written into output files.

#### Simulated RNA-seq data

In order to evaluate the accuracy and specificity of the *baerhunter*'s transcript identification and abundance estimates, a series of RNA-Seq samples were simulated using the *R* package *polyester* (Frazee *et al.*, 2015). The package creates sequencing reads given a set of genomic features and user-defined fold changes in expression between groups of samples.

1000 genomic features from *Mycobacterium tuberculosis* strain H37Rv (assembly ASM19595v2), including 24 annotated small RNAs not overlapping coding regions in the original annotation, were selected. A subset of 200 genes had artificial UTRs added to them (all 100 nucleotides long). Of these 200 genes, 50 genes were extended in the 5' end and 50 were extended in the 3' end by elongating the gene in either direction. The other half of the genes had the UTRs added as separate transcripts so that the simulation software would mimic a situation (common in bacterial expression) where the UTR and the CDS display a very different coverage in reads. Sequences in FASTA format were extracted for the final set of genes, UTRs and small RNAs and were used to produce simulated RNA-seq reads using the package *polyester*.

The package *polyester* allows the user to define fold changes between two conditions for each transcript. We provided polyester with a matrix where 10% of the features were upregulated and 10% were downregulated in one condition (i.e. 200 features were differentially expressed). Fold changes between 1 and 5 were applied.

Importantly, the current version of *polyester* does not appear to support strand-specific simulations; therefore, the development version code with minor alteration was loaded directly into the workspace to allow the production of strand-specific reads. Both single-end and paired end experiment simulations were carried out (results are reported for paired-end data only but single-end results were similar). Two conditions with 5 replicates in each condition were simulated. Every sample had 20x coverage, with other parameters set to default, e.g. read length of 100 nt.

The reads output from the *polyester* simulation were mapped to the *Mtb* H37Rv genome using the Rsubread package. The resulting BAM files were sorted with Rsamtools (Morgan *et al.*, 2019) to make them suitable for visualisation in Artemis (Carver *et al.*, 2012).

#### Real RNA-seq data

To assess the performance of *baerhunter* on real RNA-seq data, the study of (Cortes *et al.*, 2013) was used. This study was selected because it includes transcription start-site data for starved and exponentially grown cultures of *Mtb*, which can be used to benchmark predictions of non-coding RNAs and UTRs. The raw data (FASTQ files) were downloaded from ArrayExpress (dataset accession: E-MTAB-1616). A total of 6 samples were examined: ERR262980, ERR262982, and ERR262983, obtained from exponential cultures, and ERR262984, ERR262987 and ERR262988 from starved cultures. The adapters were removed, using Trimmomatic 0.32 (Bolger *et al.*, 2014) and reads were trimmed of polyA tails (added during sequencing) using fqtrim v.0.94 (Pertea); then the reads were mapped to the *Mtb* H37Rv genome (assembly ASM19595v2, GenBank accession AL123456.3) using the BWA aligner (Li and Durbin, 2009).

The resulting BAM files were used as input for the *baerhunter* analysis; several combinations of coverage cut-off values were examined and results from four combinations are included in Supplementary Table 2: 2 and 10, 5 and 10, 2 and 20 and 5 and 20. The resulting GFF3 files were used for transcript abundance quantification and differential expression analysis. The sRNA predictions were compared to a set of experimentally confirmed small RNAs provided in a review of sRNA in mycobacteria (Haning *et al.*, 2014).

#### Processing simulated and real RNA-seq data with Rockhopper

Rockhopper (downloaded from: https://cs.wellesley.edu/~btjaden/Rockhopper/) was provided with the same reference genomic sequence that was used for all *baerhunter* runs. Reads for the simulated runs were provided in FASTQ format as pairs of files holding the mate reads and the strand-specific protocol was selected.  Reads for the real RNA-seq data were provided in BAM files (mapped by us, as described in Supplementary Methods) but Rockhopper remaps these anyway using their own code. A PTT-formatted protein table (required by Rockhopper) was built using feature tables from the NCBI genome repository and is available as part of the data that can be downloaded to reproduce this analysis.

### Supplementary Table 1

ncRNA features in the simulated dataset as annotated by *baerhunter* and Rockhopper

**sRNA predictions (true number of sRNA = 24)**

| Method | All sRNA | Exact matches |
|---|---|---|
| bh_5-10 | 24 | 24 |
| bh_5-20 | 24 | 24 |
| bh_2-10 | 24 | 24 |
| bh_2-20 | 24 | 24 |
| rh_0.2-40 | 24 | 23 |
| rh_0.5-40 | 23 | 22 |

**UTR predictions (true number of UTRs = 200)**

| Method | All UTRs | Length=100 | 80<=Length<100 | Length < 80 |
|---|---|---|---|---|
| bh_5-10 | 200 | 110 | 90 | 0 |
| bh_5-20 | 200 | 110 | 90 | 0 |
| bh_2-10 | 201 | 167 | 33 | 1 |
| bh_2-20 | 200 | 167 | 33 | 0 |
| rh_0.2-40 | 199 | 100 | 62 | 36 |
| rh_0.5-40 | 196 | 96 | 52 | 47 |

Methods in the table are:
bh = baerhunter
bh_he = baerhunter results filtered to keep "high expression" features only
rh = Rockhopper
5-10, 5-20, 2-10, 2-20 are combinations of parameters for baerhunter
0.2-40 and 0.5-40 are combinations of parameters for Rockhopper (minimum expression for ncRNAs and minimum transcript length).

### Supplementary Table 2

ncRNA features in the Cortes *et al.* dataset, annotated by *baerhunter* and Rockhopper and their TSS/literature support

##### How many predicted ncRNAs are supported by TSS data

###### sRNA predictions

| Method | All sRNAs | sRNA_TSS.1 | sRNA_TSS.5 | sRNA_TSS.10 | sRNA_TSS.20 |
|---|---|---|---|---|---|
| bh_5-10 | 1439 | 913 | 979 | 998 | 1012 |
| bh_5-20 | 706 | 524 | 570 | 579 | 589 |
| bh_2-10 | 1163 | 551 | 690 | 736 | 784 |
| bh_2-20 | 586 | 292 | 393 | 427 | 452 |
| bh_he_5-10 | 235 | 192 | 209 | 212 | 214 |
| rh_0.2-40 | 568 | 303 | 341 | 359 | 393 |
| rh_0.5-40 | 309 | 155 | 189 | 212 | 232 |

###### 5’ UTR predictions

| Method | All 5’ UTRs | UTR_TSS.1 | UTR_TSS.5 | UTR_TSS.10 | UTR_TSS.20 |
|---|---|---|---|---|---|
| bh_5-10 | 605 | 251 | 289 | 316 | 340 |
| bh_5-20 | 525 | 224 | 261 | 285 | 307 |
| bh_2-10 | 912 | 192 | 274 | 329 | 372 |
| bh_2-20 | 792 | 161 | 237 | 284 | 325 |
| bh_he_5-10 | 223 | 110 | 128 | 137 | 148 |
| rh_0.2-40 | 1022 | 482 | 545 | 583 | 616 |
| rh_0.5-40 | 442 | 216 | 250 | 266 | 280 |

##### How many predicted ncRNAs are supported by data from the Haning et al. review

###### sRNA predictions

| Method | All sRNAs | sRNA_ Haning.1 | sRNA_ Haning.5 | sRNA_ Haning.10 | sRNA_ Haning.20 |
|---|---|---|---|---|---|
| bh_5-10 | 1439 | 440 | 432 | 422 | 404 |
| bh_5-20 | 706 | 301 | 298 | 292 | 281 |
| bh_2-10 | 1163 | 398 | 388 | 383 | 375 |
| bh_2-20 | 586 | 268 | 263 | 263 | 256 |
| bh_he_5-10 | 235 | 116 | 114 | 112 | 109 |
| rh_0.2-40 | 568 | 268 | 259 | 254 | 219 |
| rh_0.5-40 | 309 | 167 | 165 | 160 | 142 |

###### UTR predictions

| Method | All UTRs | UTR_ Haning.1 | UTR_ Haning.5 | UTR_ Haning.10 | UTR_ Haning.20 |
|---|---|---|---|---|---|
| bh_5-10 | 975 | 383 | 377 | 369 | 354 |
| bh_5-20 | 815 | 354 | 349 | 342 | 331 |
| bh_2-10 | 1405 | 488 | 480 | 471 | 451 |
| bh_2-20 | 1144 | 445 | 435 | 428 | 413 |
| bh_he_5-10 | 280 | 164 | 163 | 160 | 156 |
| rh_0.2-40 | 1662 | 332 | 309 | 279 | 234 |
| rh_0.5-40 | 676 | 198 | 189 | 168 | 139 |

The total number of ncRNA features (“All sRNAs” and “All UTRs”) annotated by baerhunter and Rockhopper methods run with different parameters using RNA-seq data from exponentially grown and starving cultures of *Mtb* (Cortes *et al.*, 2013) are presented together with the subset of these features that are backed up by external evidence (TSS and literature).

Methods in the table are:

bh = baerhunter

bh_he = baerhunter results filtered to keep “high expression” features only

rh = Rockhopper

5-10, 5-20, 2-10, 2-20 are combinations of parameters for baerhunter

0.2-40 and 0.5-40 are combinations of parameters for Rockhopper (minimum expression for ncRNAs and minimum transcript length).

The columns labeled TSS.x correspond to the number of non-coding RNAs that are within x nucleotides of a TSS listed in the Cortes *et al.* study.

The columns labeled Haning.x refer to ncRNAs that have at least x nucleotides overlap with a ncRNA listed in the Haning *et al*. review (the number includes both “confirmed” and “unconfirmed” ncRNAs).

#### Additional Supplementary Figures

#### Supplementary Figure 4

Length distributions for UTRs predicted in the simulated RNA-seq data

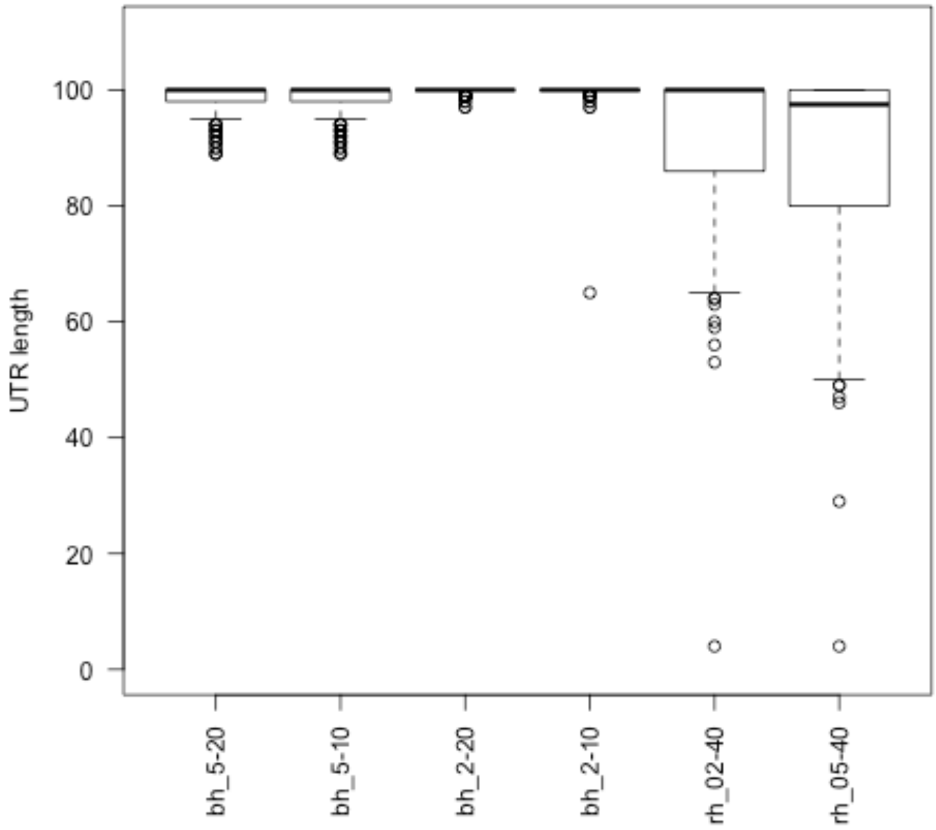


Box-and-whisker plots representing the lengths of UTRs predicted by *baerhunter* and Rockhopper runs (with different parameters) in the simulated RNA-seq data. 200 UTRs of length 100 were added to the simulation. The reads are simulated in a way that adds noise to the signal – this explains why the programs do not always return UTRs of length 100.

#### Supplementary Figure 5

Baerhunter and Rockhopper predictions of sRNAs in the Cortes *et al.* RNA-seq dataset and their TSS support

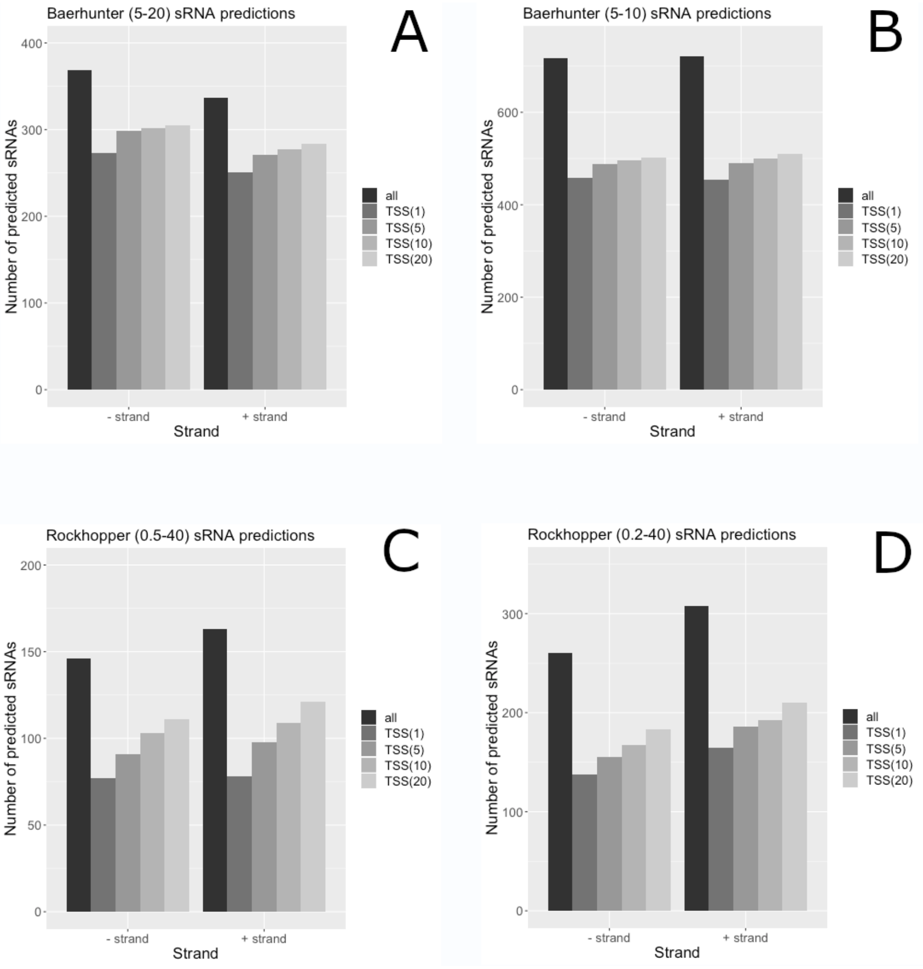


The total number of predicted sRNAs (black) is contrasted with the number of predictions supported by an experimentally detected TSS at different resolution cut-offs (grey bars). TSS(x) means that an experimentally derived TSS is found within x nucleotides of the predicted sRNA's 5' end.

A) and B) Baerhunter predictions; C) and D) Rockhopper predictions.

#### Supplementary Figure 6

Predicted features of weak expression are often supported by weak TSS signal

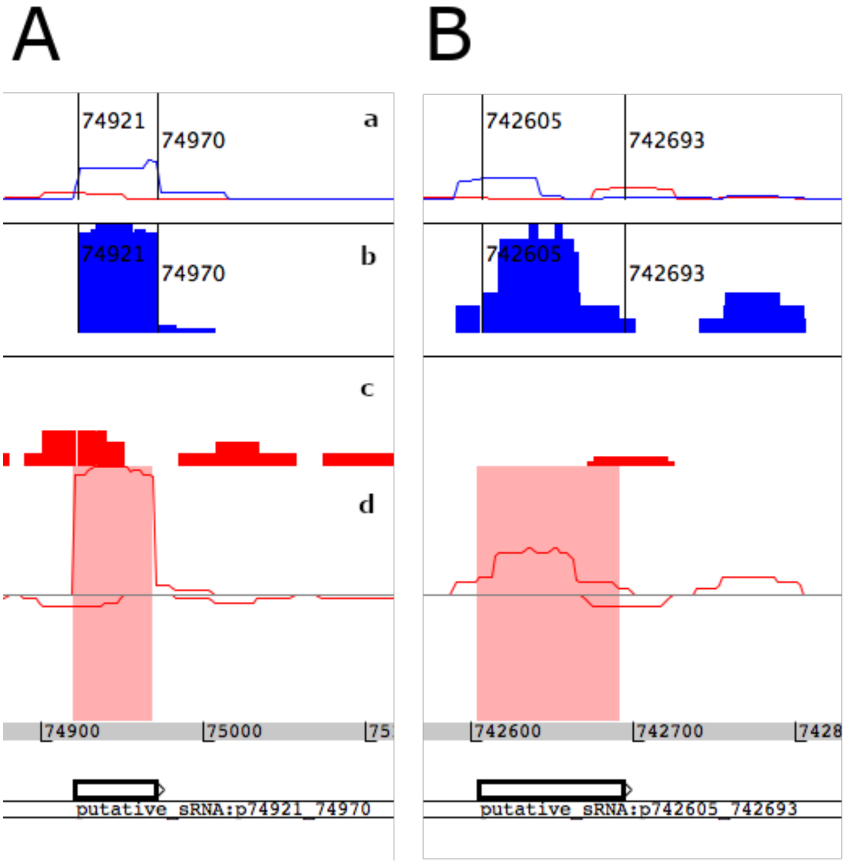


Due partly to its naïve approach for identifying expressed regions, *baerhunter* will pick up areas of expression in the RNA-seq signal that do not correspond to a reported TSS or known ncRNA. Occasionally, these features will correspond to a region that shows clear (albeit usually low) expression and their presence is supported by a weak TSS, as shown in both examples in this figure. Rows correspond to a) TSS from the Cortes et al. study; b and c) coverage files produced by Rockhopper for the positive (blue) and negative (red) strand; and d) coverage from our mapping of the RNA-seq data to the reference genome. Predicted sRNAs are highlighted in pink rectangles and their coordinates are marked by the vertical lines in rows (a) and (b).

#### Supplementary Figure 7

Baerhunter and Rockhopper predictions of 5' UTRs in the *Cortes et al.* RNA-seq dataset and their TSS support

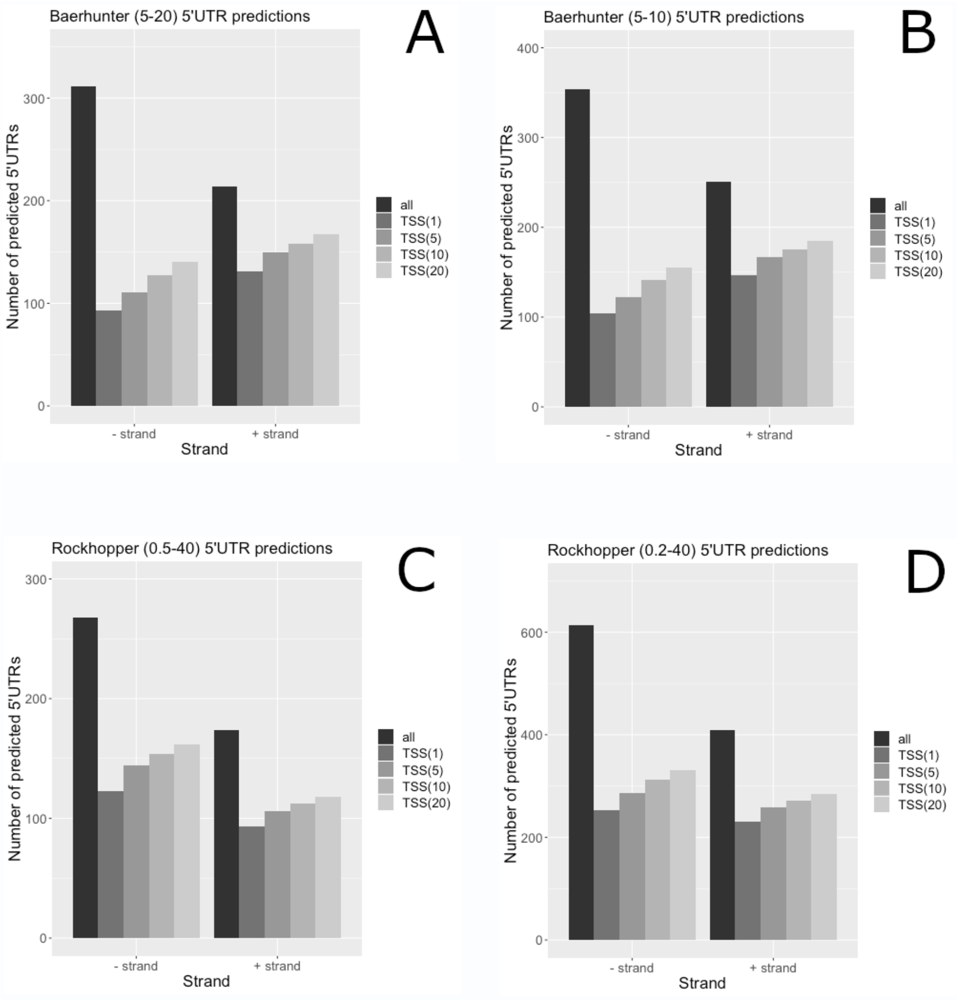


The total number of predicted 5' UTRs (black) is contrasted with the number of those predicted 5' UTRs that are also supported by an experimentally detected TSS at different resolution cut-offs (grey bars). TSS(x) means that an experimentally derived TSS is found within x nucleotides of the predicted UTR's 5' end. UTRs predicted to span the space between two genes so that they are both 3' to one CDS and 5' to another, have not been included as it is not possible to assign them with certainty to the upstream or downstream gene; these expressed regions are part of the intergenic space in operons and many of them would not have TSS associated with them anyway. A & B) *baerhunter* predictions; C & D) Rockhopper predictions.
